# Analysis of intratumoral immune heterogeneity reveals spatial organisation of immunosuppression in breast cancer

**DOI:** 10.64898/2025.12.18.694622

**Authors:** Christina Metoikidou, Pierre-Emmanuel Bonté, Vadim Karnaukhov, Laetitia Lesage, Lounes Djerroudi, André Nicolas, Philemon Sirven, Anne Vincent-Salomon, Olivier Lantz, Emanuela Romano, Sebastian Amigorena

## Abstract

Recent advances in single-cell transcriptomics have illuminated intratumoral heterogeneity (ITH) in breast cancer, yet the spatial organization and functional states of immune populations within luminal tumors remain poorly understood. Here, we generated a spatially and clonally resolved single-cell immune atlas of early-stage luminal breast cancer by integrating paired single-cell RNA/TCR sequencing with multiplex immunohistochemistry across three spatially distinct tumor regions: central (core-proximal), intermediate and peripheral (invasive margin-proximal), in five patients. Peripheral regions are enriched in naïve CD4 T cells, monocytes, dendritic cells, and memory CD8 T cells with bystander properties, alongside TCF1^+^ exhausted progenitors (Tpex), and displayed elevated antigen presentation and tertiary lymphoid structures (TLS) signatures. In contrast, central regions harbor macrophages, clonally expanded regulatory T cells (Tregs), and two subsets of terminally exhausted CD8 T cells (Tex), one tissue-resident and one circulating, both expressing tumor-reactive features. Clonal overlap was observed between peripheral Tpex and central, tissue-resident Tex cells, revealing a spatially confined differentiation trajectory toward exhaustion within the tumor microenvironment. Central T cells also upregulate TGF-β signaling and metabolic pathways including fatty acid oxidation and oxidative phosphorylation, consistent with adaptation to an immunosuppressive microenvironment. Together, these findings reveal a spatially structured immune landscape in luminal breast cancer, marked by progressive immunosuppression from the tumor margin to the core, providing a framework for future region-specific immunotherapeutic strategies.

## INTRODUCTION

Breast cancer (BC) is the most common malignancy among women worldwide (*1*) and is clinically stratified into subtypes based on estrogen receptor (ER), progesterone receptor (PR), and human epidermal growth factor receptor 2 (HER2) status (*2*). Luminal breast cancers (LBCs), which are ER-positive, comprise luminal A (ER^+^, PR^+/-^, HER2^-^) and luminal B (ER^+^, PR^+/-^, HER2^+/-^) subtypes (*3*). Compared with triple-negative and HER-2 enriched tumors, which are immunologically active, luminal tumors are often immune-quiescent, characterized by limited immunogenicity and fewer therapeutic options (*3*).

Despite their relatively low immune activity, luminal tumors account for the majority of breast cancer cases (*1*), yet the composition, functional states, and spatial organization of their tumor microenvironment (TME) remain poorly characterized, limiting opportunities to identify prognostic markers or therapeutic targets. Of note, recent studies have begun exploring immunotherapeutic strategies in hormone receptor-positive tumors, underscoring a growing interest in activating antitumor immunity within this previously unresponsive subtype (*4, 5*). These efforts highlight the need for detailed mapping of the luminal TME, including the spatial distribution and functional states of immune cells, to better understand mechanisms of immune invasion and guide potential therapeutic interventions.

Prognostic and predictive biomarkers, such as tumor-infiltrating lymphocytes (TILs) (*6*), PD-L1 expression (*7*), and tumor mutational burden (TMB) (*8, 9*) provide insights into immune engagement (*10*), but often fail to reliably predict clinical outcomes. A key limitation is that biomarker assessment relies on small tissue biopsies, which, due to their size, may not fully capture the complexity of the breast TME (*11*). *In situ* biopsies are generally difficult to direct precisely to specific regions of the tumors, introducing possible sampling biases, with, for example, the spatial positioning of T cells at the invasive margins versus the tumor core influencing prognostic significance (*12*).

While the overall presence of TILs correlates with better prognosis (*13*), the functional state and spatial distribution of T cells add other layers of complexity. CD8 T cells are central to antitumor immunity (*14*), but undergo exhaustion in the TME (*15*), progressing from progenitor, memory-like states (Tpex) to terminally exhausted (Tex) states (*16*). Tpex cells retain stem- and memory-like traits, sustaining long-term responses (*17–19*). Recent studies from our group and others, identified distinct subsets of Tpex, originating either from circulating lymphocytes in peripheral blood, or from tissue-resident memory T cells (*20, 21*). Both Tpex subsets differentiate intratumorally into Tex cells (*16, 20, 21*). While Tex cells are considered functionally impaired, emerging evidence suggests that they undergo clonal expansion and proliferation, retain tumor-reactive properties (*22–24*) and can be associated with favorable prognosis and response to immune checkpoint blockade (ICB) therapies (*24–26*). How Tpex and Tex are spatially distributed within luminal tumors and how their clonal dynamics vary across regions remains, however, largely unknown.

Beyond CD8 T cells, other immune populations also shape the TME. Regulatory T cells (Tregs) and macrophages promote immunosuppression (*27–29*), whereas monocytes and dendritic cells (DCs) support immune surveillance and tertiary lymphoid structure (TLS) formation (*30, 31*). Although single-cell transcriptomic analyses have characterized immune heterogeneity, the spatial organization of these populations in luminal tumors remains poorly defined.

To address this gap, we performed high-resolution profiling of immune cells across three distinct anatomical regions of luminal breast tumors: central (proximal to the tumor core), intermediate and peripheral (proximal to the invasive margins), using single-cell RNA/ TCR sequencing with feature barcoding and multiplex immunohistochemistry (IHC). This approach enabled detailed characterization of immune cellular states, clonality and transcriptomic profiling, while retaining coarse spatial context. Peripheral regions are enriched in monocytes, DCs, naïve CD4 and memory-like Tpex CD8 T cells, aligning with immune surveillance and TLS activity. In contrast, central tumor regions harbor macrophages, clonally expanded Tregs and Tex CD8 T cells with tumor-reactive properties. Clonal analysis revealed that central tissue-resident Tex share clones with peripheral Tpex cells, suggesting a spatial gradient of intratumor CD8 T cell differentiation. Furthermore, T cells within central regions upregulate TGF-β signaling, fatty acid (FA) metabolism, and oxidative phosphorylation (OXPHOS) signatures, reflecting metabolic adaptation to an immunosuppressive TME. Collectively, these findings reveal a compartmentalized immune landscape in luminal breast tumors and highlight how spatial context shapes immune composition, functional states, and clonal dynamics, providing a framework for future region-targeted therapeutic strategies.

## RESULTS

### Patients and study design

To investigate the spatial organization of immune cell subsets in the TME, we profiled tumors from five patients with early-stage, untreated luminal breast cancer (LBC), all diagnosed with invasive ductal carcinoma (IDC) (Figure 1A and Supplemental Table 1). Two patients were diagnosed with luminal A (P1 and P2) and three with luminal B BC type (P3-P5) (Supplemental Table 1). Following surgery, a pathologist collected three fresh tissue samples per patient, each from a macroscopically distinct tumor region: one proximal to the tumor core, one proximal to the invasive margins and one intermediate localized between these two regions (Figure 1A and Supplemental Figure 1A). This approach allowed us to determine whether each biopsy captures a different immune microenvironment and contributes to intratumoral heterogeneity. All samples underwent hashing antibody labeling, followed by single-cell RNA- and TCR-sequencing (Figure 1A).

**Figure 1.**
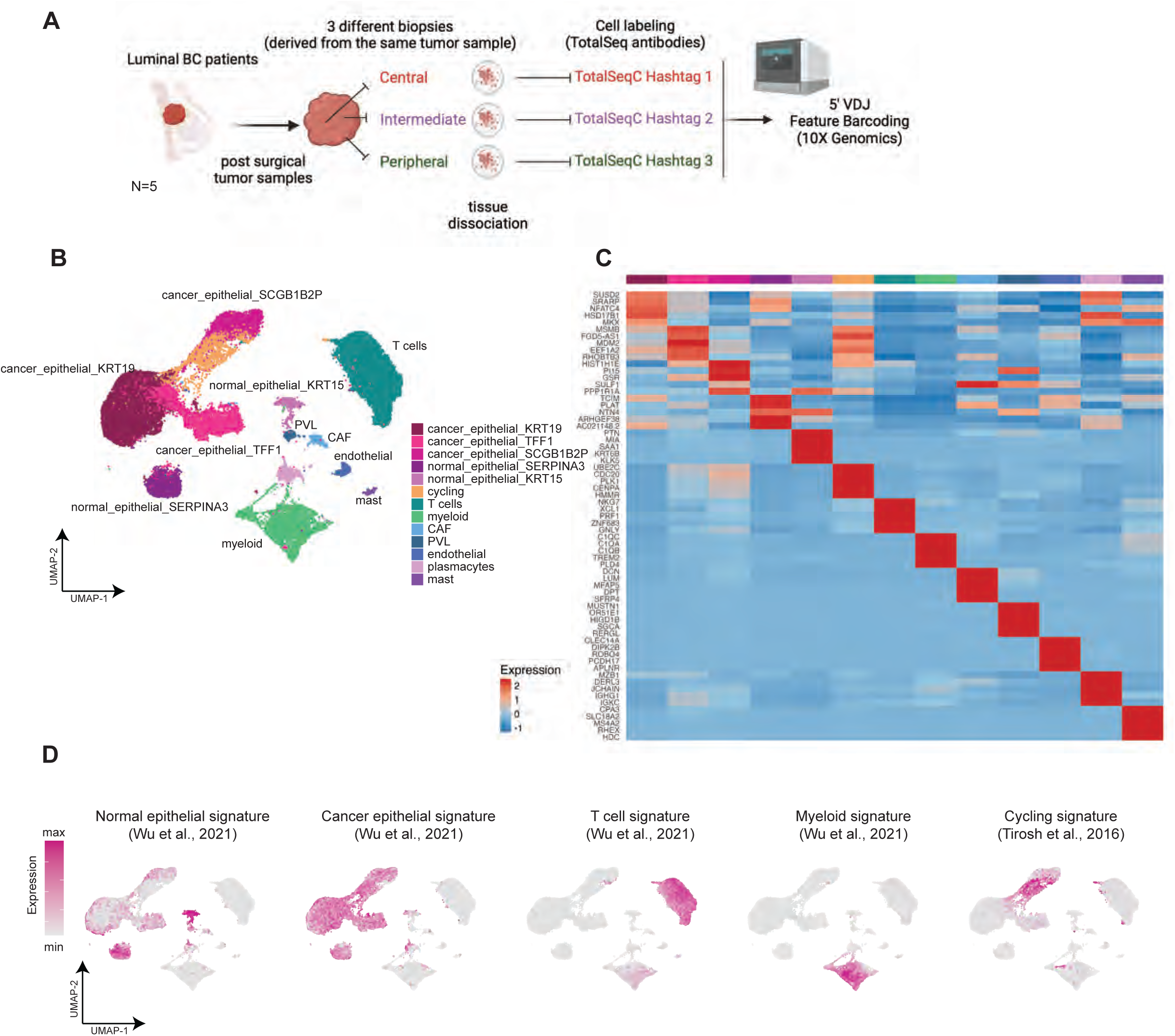
Study design and spatial profiling of immune and non-immune cell types in luminal breast cancer. **(A)** Graphical overview of the study design. Workflow for sample collection and processing for scRNA/TCR sequencing. Three distinct post-surgical tumor-tissue samples were obtained for each patient, derived from tumor regions proximal to tumor-core, intermediate and invasive margin respectively. A total of 15 samples (from 5 patients, 3 samples from each patient) were analyzed. Samples were labelled with antibody hashtags and sequenced using 5’ V(D)J 10X technology with Feature Barcoding. Created with BioRender.com. **(B)** UMAP of cell clusters obtained from 52,812 tumor-infiltrating single cells. CAF: cancer-associated fibroblast cell; PVL: perivascular cell. **(C)** Heatmap of normalized expression of top5 differential expressed genes (DEGs) per cluster (wilcoxon rank sum test). **(D)** Feature plot showing the normalized expression of gene signatures in tumor-infiltrating cells.

### Distinct transcriptomic landscape across tumor-regions in luminal BC

For all samples, we collected information on both transcriptomic profiles and tumor region of origin (central, intermediate or periphery) through hashtag labeling. After quality control and filtering (see “Methods”), we selected a total of 52,812 single cells. Graph-based clustering of all cells resulted in 13 clusters, including five of epithelial origin, as well as clusters of T lymphocytes, myeloid, plasma, mast, endothelial, perivascular (PVL) cells and cancer associated fibroblasts (CAF) (Fig. 1B). Differential gene expression (DE) analysis (Figure 1C and Supplemental Table 2) and projection of a selected set of breast cancer gene signatures (*32, 33*), defining cell states, revealed the identity of the clusters (Figure 1D, Supplemental Figure 1B and Supplemental Table 3). Among the five epithelial clusters, two upregulated normal epithelial markers (normal_epithelial_SERPINA3 and normal_epithelial_KRT15), and three overexpressed cancer epithelial gene signatures (cancer_epithelial_KRT19, cancer_epithelial_TFF1 and cancer_epithelial_SCGB1B2P) (Figure 1D). Cancer epithelial cells and T lymphocytes were the most abundant populations (Supplemental Figure 1C).

All clusters were represented across all patients, with cancer_epithelial_SCGB1B2P cells being enriched in luminal B samples (P3-P5) (Supplemental Figure 1, D-E). In single cell studies, cycling cells tend to cluster together, due to upregulation of genes involved in cell cycling, independently of their cell state of origin. To identify the origin of cycling cells, we performed label transfer from the non-cycling cells onto the cycling subset (Supplemental Figure 1F). This analysis unveiled that most cycling cells are cancer epithelial cells, followed by T cells (Supplemental Figure 1F).

To investigate regional transcriptomic landscapes, we compared cell cluster abundancies across regions. For each patient, three spatially distinct samples were tagged with unique hashtag antibodies and pooled together for sequencing. To assign the samples back to their respective tumor region of origin we performed hashtag demultiplexing (Supplemental Figure 2, A-D). This analysis unveiled cells labelled by one hashtag and therefore uniquely assigned to one tumor region (singletons), as well as cells either labeled with two hashtags (doublets), or exhibiting very low hashtag expression (negative) (Supplemental Figure 2A), precluding their unique assignment to a tumor region. Downstream analysis for comparisons across tumor regions was performed on singletons.

To map spatial organization within tumors, we visualized cell state densities across central, intermediate and peripheral regions (Figure 2A). This analysis revealed distinct regional distributions, further supported by hierarchical clustering of cell state frequencies, which grouped samples by spatial origin (Figure 2B), indicating a conserved spatial organization of the TME. The tumor core is mainly populated with cancer epithelial cells (epithelial_KRT19), while samples from regions proximal to the invasive margins contain more normal epithelial cells (epithelial_SERPINA3) and T-cells (Figure 2C). These results were consistent across all patients (Fig. 2D), with no significant regional differences in other clusters (Supplemental Figure 2, E-F).

**Figure 2.**
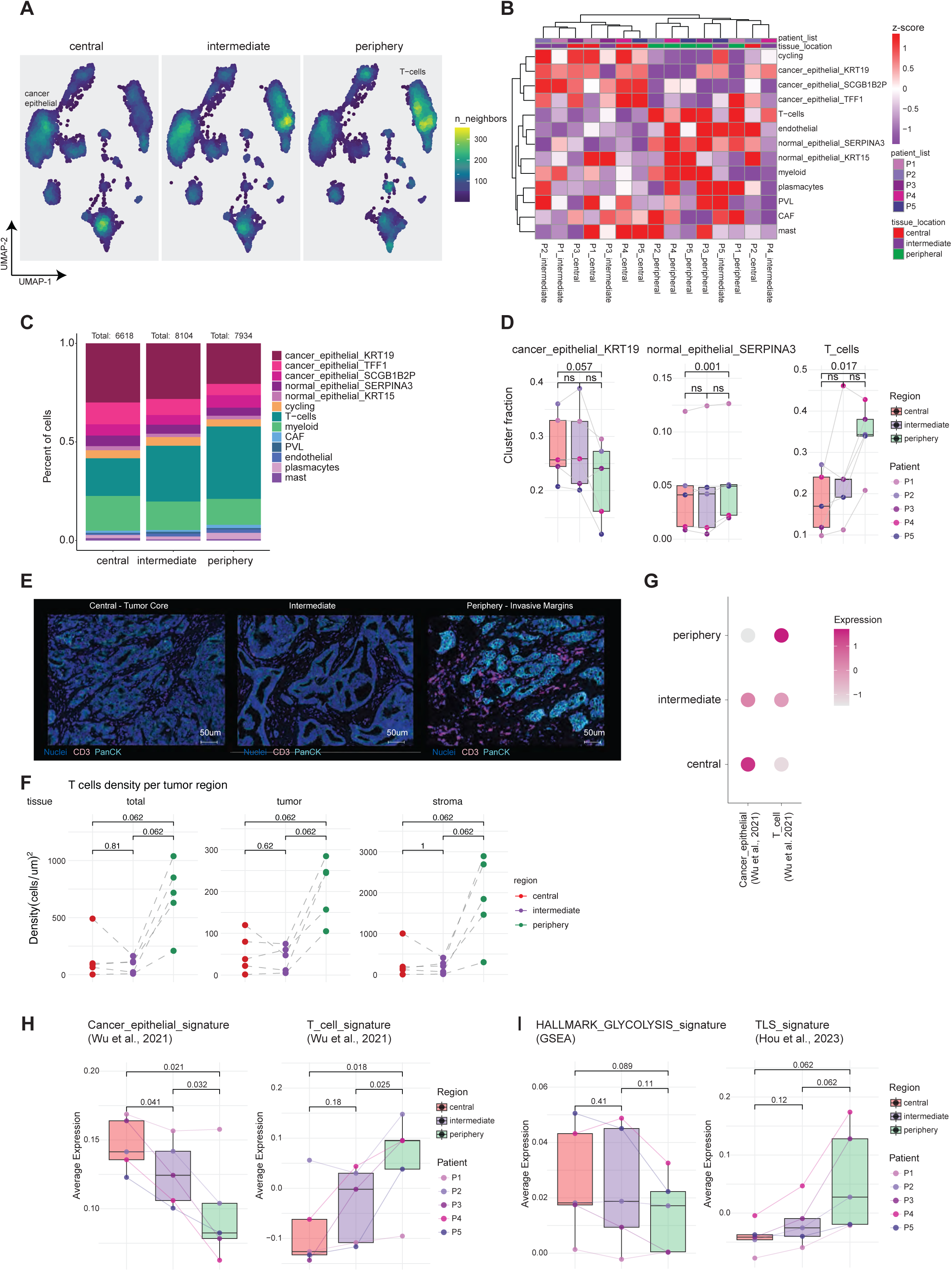
Identification of distinct spatial patterns of cell type distribution across tumor regions. **(A)** UMAP of cell states density across tumor regions: central (proximal to tumor core), periphery (proximal to invasive margins) and intermediate. Data include n=5 samples per region, totaling n=15 samples from N=5 patients. Analysis performed on singlet cells based on HTO expression. **(B)** Unsupervised hierarchical clustering heatmap depicting the normalized distribution of cell clusters’ fractions across tumor regions in individual patients. Analysis performed on singlet cells based on HTO expression. Data were first normalized within each patient across tumor regions, and then z-scored across all patients and samples (by row). Clustering was performed using Euclidean distance to group both rows (representing cell clusters) and columns (representing tumor regions and patient samples). Clustering reveals patterns of cell states’ distribution based on spatial location within the tumor. **(C)** Bar plot showing the relative abundance of cell clusters per tumor-region. Analysis performed on singlet cells based on HTO expression. Data include n=5 samples per region, totaling n=15 samples from N=5 patients. The absolute number of cells per tumor region is indicated above each bar. **(D)** Cluster frequencies per tumor region. Analysis performed on singlet cells based on HTO expression. Each dot represents a single sample (n = 15). Samples were derived from N = 5 patients, with 3 tumor-region samples (central, intermediate, and periphery) collected from each patient. Individual patients are represented by colored dots, and lines connect samples from the same patient. Statistical comparisons between regions were conducted using a two-way repeated measures ANOVA with Greenhouse-Geisser correction. Tukey’s post-hoc test was applied for pairwise comparisons. Adjusted p-values for pairwise comparisons are indicated; ns= not significant. **(E)** Representative IHC images of CD3 T cells (in pink), nuclei (in dark blue), and pan-cytokeratin (in cyan) across tumor regions (central, intermediate, and peripheral). Scale bar: 50um. **(F)** CD3 T cell density (cells/ um^2^) across tumor regions: central-core, intermediate, and periphery-invasive margins. Data points represent individual patient samples, with each dot corresponding to a unique analysis region. Statistical comparisons were performed using the Wilcoxon signed-rank test with Bonferroni correction for multiple comparisons; p-values are indicated. **(G)** Mean expression of epithelial cancer cells- and T cells-gene signatures per tumor-tissue region. Analysis performed on singlet cells based on HTO expression. **(H)** Boxplots of cancer epithelial cells (left panel) and T cells (right panel) gene signatures expression across tumor regions. Analysis performed on singlet cells based on HTO expression. Points represent individual samples from N=5 patients, with lines connecting paired samples from the same patient across regions. Statistical comparisons were performed using paired t-tests with significance indicated by p-values for pairwise comparisons: central vs. intermediate, intermediate vs. periphery, and central vs. periphery. **(I)** (left panel): Expression of the HALLMARK_GLYCOLYSIS signature across tumor regions. Pairwise comparisons were conducted using the t-test (paired); (right panel): Expression of the TLS signature across different tumor regions (central, intermediate, and periphery) for each patient. Pairwise comparisons were performed using the Wilcoxon test (paired); Analysis performed on singlet cells based on hashtag-antibody expression. Points represent individual samples from N=5 patients, with lines connecting paired samples from the same patient across regions. Statistical tests were chosen based on the normality of the data, with non-parametric tests applied for non-normally distributed data and parametric tests for normally distributed data (as assessed by Shapiro-Wilk).

To validate these findings, we performed multiplex IHC on matched tissue sections (Supplemental Figure 2G) confirming higher CD3^+^ T cell density in peripheral regions compared to the core (Figure 2, E and F) and greater enrichment in stromal versus tumor compartments (Supplemental Figure 2H).

To assess functional states, independently of clustering, we analyzed gene signature expression (*32*) across tumor areas. Central regions upregulated cancer epithelial and glycolytic programs (Figure 2, G-I), particularly within cancer epithelial cells and CAFs (fig. S2I), indicating a metabolically adapted suppressive niche. Peripheral regions upregulated T-cell, TLS (*34*) and antigen-presentation signatures (Figure 2, G-I, and Supplemental Figure 2, I and J), mainly in T cell and myeloid clusters (Supplemental Figure 2I). This pattern coincided with enrichment of B cells and antigen presenting cells (APCs) in TLS-rich zones, supporting their role in local immune activation. Consistently, B cell (*32*) and antigen-processing gene signatures (table S3) were elevated in the periphery, linking TLS formation to enhanced immune surveillance at these regions (Supplemental Figure 2J). Together, these data reveal an immune-metabolic compartmentalization, with the tumor core maintaining a glycolytic, suppressive state and the periphery fostering antigen presentation and adaptive immunity.

### Myeloid and B cell states across tumor regions

Next, we examined the distribution and functional states of myeloid subsets across tumor regions. To do so, we extracted myeloid cells and reanalyzed them independently of other cell types, identifying 3,999 myeloid cells grouped into 12 clusters (Figure 3A). DE analysis and projection of myeloid state markers (*35*) (Supplemental Figure 3, A-D and Supplemental Table 4) revealed six macrophage clusters, two DC clusters, and monocyte subsets, encompassing both classical (CD14^high^) and non-classical (*FCGR3A*/CD16^high^) types (Supplemental Figure 3D), along with plasmacytoid DCs (pDCs), cycling myeloid cells, and a small B-cell subset, that co-clustered with myeloid populations in the global UMAP.

**Figure 3.**
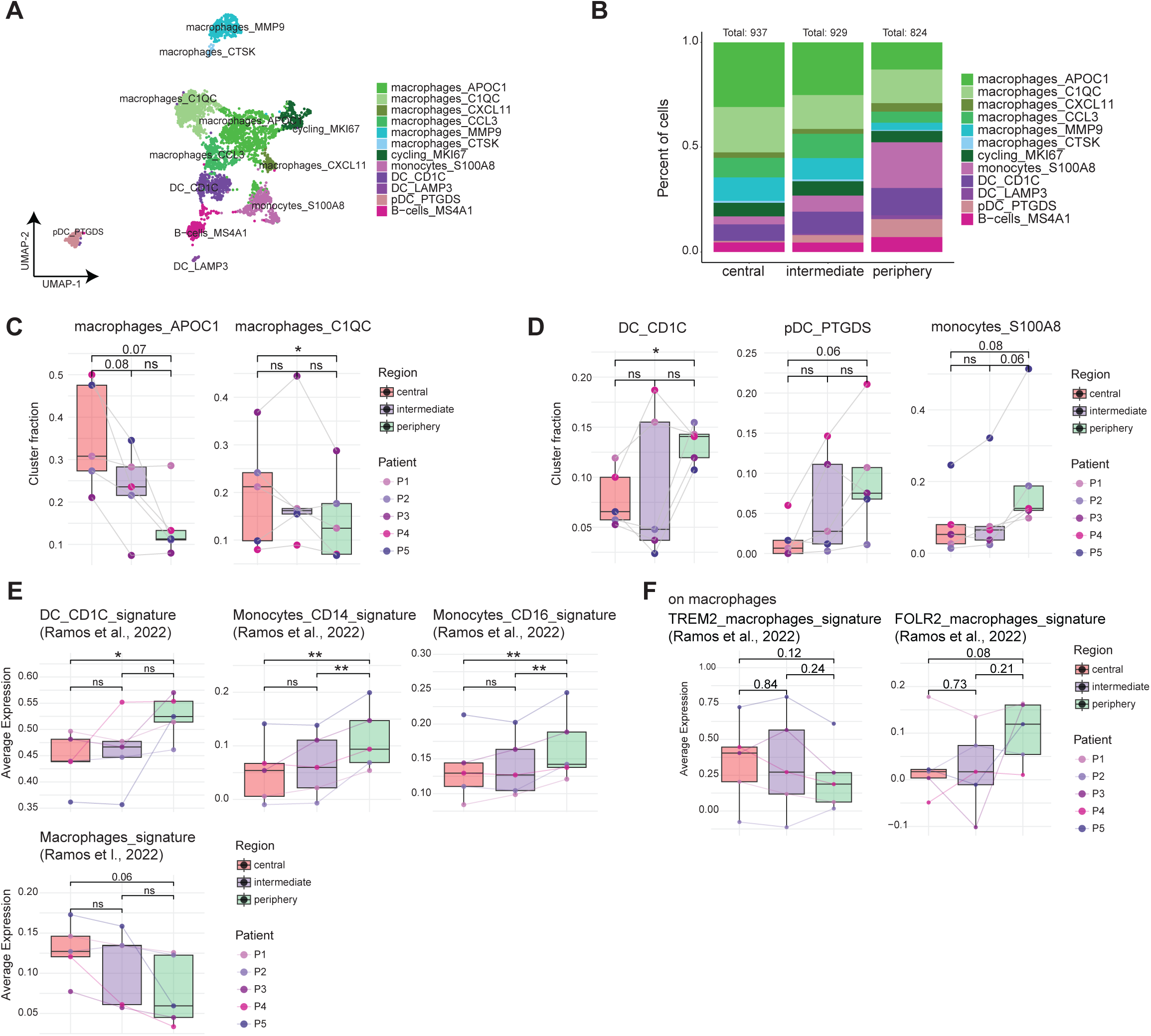
Myeloid and B cell states across tumor-tissue regions in luminal breast cancer. **(A)** UMAP of 3,999 single tumor-infiltrating myeloid cells. **(B)** Relative abundance of myeloid clusters per tumor-region. Data include n=5 samples per region (central, intermediate, periphery), totaling n=15 samples from N=5 patients. Analysis performed on singlet myeloid cells based on HTO expression. Absolute cell number is indicated above bars. **(C-D)** Cluster frequencies per tumor region. Each dot represents a single sample (n = 15). Samples were derived from N = 5 patients, with 3 tumor-region samples (central, intermediate, and periphery) collected from each patient. Individual patients are represented by colored dots; lines connect matched samples. Analysis performed on singlet myeloid cells based on HTO expression. Statistical comparisons between tumor regions were conducted using a two-way repeated measures ANOVA with Greenhouse-Geisser correction. Tukey’s post-hoc test was applied for pairwise comparisons. Adjusted p-values for pairwise comparisons are indicated; p ≤ 0.05 (*); ns= not significant. **(E)** Expression of gene signatures related to myeloid cell state across tumor regions. Analysis performed on singlet myeloid cells based on HTO expression. Pairwise comparisons were conducted using the t-test (paired). Points represent individual samples from N=5 patients; lines connect matched samples. Adjusted p-values for pairwise comparisons are indicated; p ≤ 0.05 (*); p ≤ 0.01 (**); ns= not significant. **(F)** Expression of gene signatures related to macrophages state across tumor regions. Analysis performed on subsetted macrophage cells (including macrophages_APOC1, _C1QC, _CXCL11, _CCL3, _MMP9, _CTSK). Pairwise comparisons were conducted using the t-test (paired). Points represent individual samples from N=5 patients, with lines connecting paired samples from the same patient across regions. Adjusted p-values for pairwise comparisons are indicated.

Among macrophage subsets, two clusters (macrophages_APOC1 and macrophages_C1QC) upregulated M2-like/ tumor-associated macrophages (TAMs) genes, including lipid-associated and immunomodulatory markers such as *APOC1*, *GPNMB* and *FABP5*, while two clusters (macrophages_CXCL11 and macrophages_CCL3) upregulated pro-inflammatory M1-like genes, including *APOBEC3A, CXCL11* and *IL1B* (Supplemental Figure 3A). Two additional clusters (macrophages_MMP9 and macrophages_CTSK) expressed matrix metalloproteinases (MMPs), consistent with roles in extracellular matrix remodeling (Supplemental Figure 3A). M2-like macrophages represented the dominant myeloid populations (Supplemental Figure 3E), with all myeloid subsets present across patients (Supplemental Figure 3F).

To determine their spatial organization, we compared myeloid subset abundancies across tumor regions. M2-like macrophages were enriched in the tumor core (Figure 3, B and C, and (Supplemental Figure 3G), whereas monocyte, pDC, and DC clusters accumulated at the invasive margins (Figure 3, B and D, and Supplemental Figure 3G). Other myeloid subsets showed no significant regional differences (Supplemental Figure 3H). To validate these findings independently of clustering, we analyzed gene signature expression patterns. Consistent with the cluster proportions analyses, monocyte and DC programs (*35*) were elevated in peripheral regions, while macrophage-associated signatures predominated in central regions (Figure 3E and Supplemental Table 3).

We next examined macrophage polarization patterns across tumor regions. FOLR2-type macrophages, associated with antigen presentation, CD8 T cell priming, and favorable prognosis (*35*), were enriched in the periphery, consistent with the increased presence of monocytes and DCs in this region (Figure 3F and Supplemental Table 3). In contrast, TREM2-type macrophages, linked to lipid metabolism, immunosuppression, and tumor progression (*36, 37*), tended to accumulate in the tumor core (Figure 3F and Supplemental Table 3).

Collectively, these results indicate region-specific functional states across the myeloid compartment, with immune-stimulatory APCs, including monocytes, DCs and FOLR2 macrophages concentrated at the invasive margins, while immunosuppressive TREM2 TAMs dominated the tumor core.

### T cell and NK cell states by tumor region

To investigate intra-tumor T cell heterogeneity, 8,853 T cells were analyzed independently, revealing 14 clusters, present across all patients (Figure 4A and Supplemental Figure 4, A and B). DE and gene signature (*21*) analysis identified distinct CD8, CD4 and NK populations (Figure 4, A-C, and Supplemental Figure 4, C-E, and Supplemental Table 5). Among CD8 T cells, CD8_Trm_CCL4 expressed memory T-cell gene signatures (e.g., CD8_IL7R_Tm) (Figure 4B and Supplemental Figure 4D), representing stem-like precursor cells (Tpex), while CD8_Tex/Trm_ZNF683 and CD8_Tex/Tcirc_GZMK exhibited late-terminal exhaustion signatures (e.g., CD8_Terminal_Tex) (Figure 4B).

**Figure 4.**
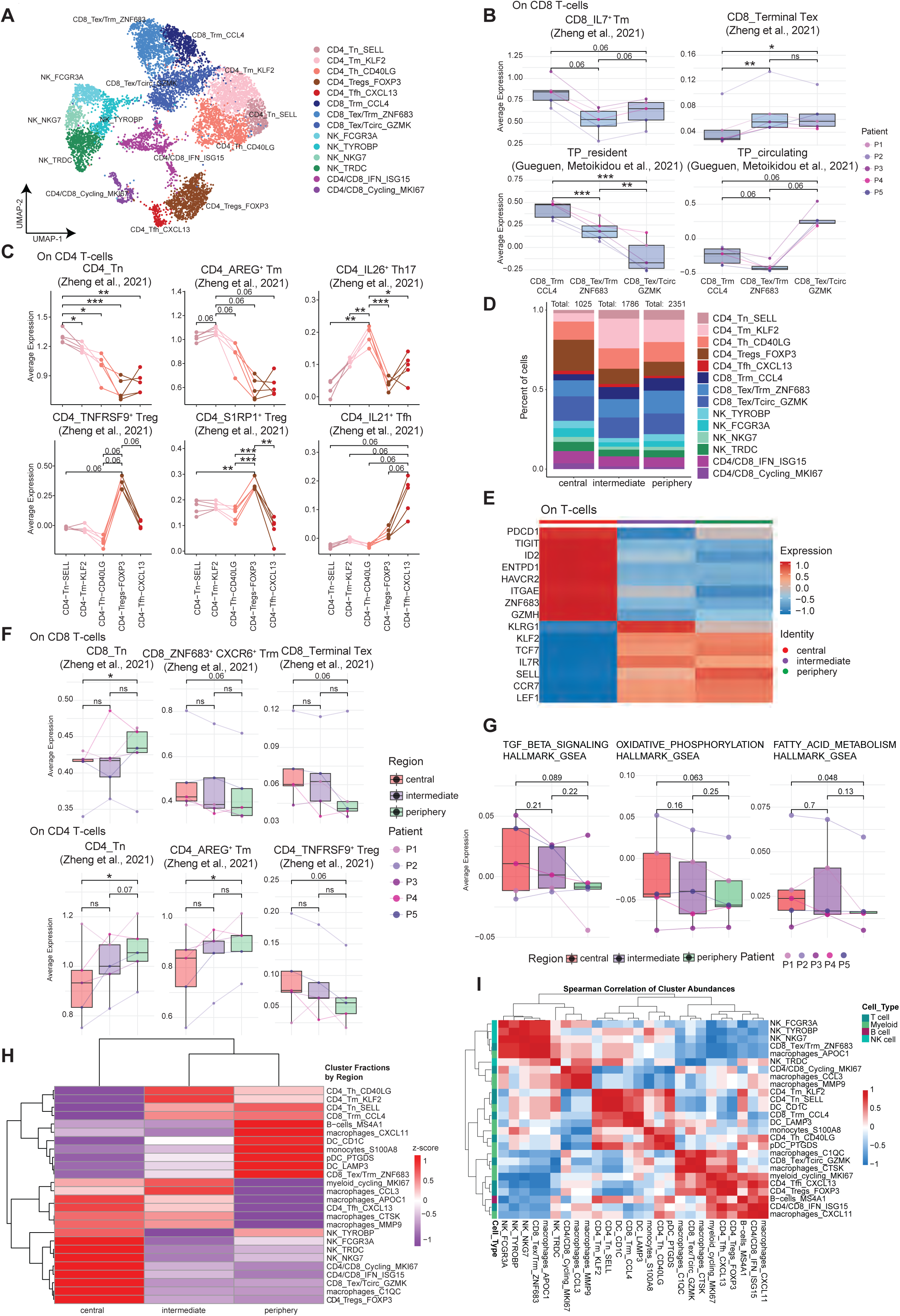
T- and NK- cell transcriptomic landscape across tumor-regions in luminal breast cancer. **(A)** UMAP of 8,853 tumor-infiltrating T-cells, color-coded by identified clusters. **(B)** Expression of memory (IL7^+^ Tm), terminal exhausted (Terminal Tex), tissue-resident (TP_resident) and circulating (TP_circulating) CD8 T-cell signatures. Analysis performed on subsetted CD8 T-cells. Points represent individual samples from N=5 patients, lines connect paired samples across regions. Pairwise comparisons were conducted using t-test (paired) or Wilcoxon test (paired) as appropriate*. Adjusted p-values for pairwise comparisons are indicated; p ≤ 0.05 (*); p ≤ 0.01 (**); p ≤ 0.001 (***); ns= not significant. **(C)** Expression of naïve (Tn), memory (AREG^+^ Tm), Th (IL26^+^ Th17), Tfh (IL21^+^ Tfh) and Treg (S1RP1^+^ Treg/ TNFRS9^+^ Treg) CD4 T-cell gene signatures. Analysis performed on subsetted CD4 T-cells. Points represent individual samples from N=5 patients, with lines connecting paired samples from the same patient across regions. Pairwise comparisons were conducted using t-test (paired) or Wilcoxon test (paired) as appropriate. Adjusted p-values for pairwise comparisons are indicated; p ≤ 0.05 (*); p ≤ 0.01 (**); p ≤ 0.001 (***); ns= not significant. **(D)** Relative abundance of T-cell clusters per tumor-region. Data include n=5 samples per region (central, intermediate, periphery), totaling n=15 samples from N=5 patients. Analysis performed on singlet T-cells based on HTO expression. Absolute cell numbers per tumor region indicated above bars. **(E)** Heatmap of averaged marker gene expression across tumor-regions. Analysis performed on singlet T-cells based on HTO expression. **(F)** (upper panel): Expression of CD8 T-cell state associated gene signatures per region. Analysis performed on subsetted CD8 T-cells. (bottom panel): Expression of CD4 T-cell state associated gene signatures per region. Analysis performed on subsetted CD4 T-cells. Points represent individual samples from N=5 patients, with lines connecting paired samples from the same patient across regions. Pairwise comparisons were conducted using the t-test (paired) or Wilcoxon test (paired) as appropriate*. Adjusted p-values for pairwise comparisons are indicated; p ≤ 0.05 (*); ns= not significant. **(G)** Expression of T-cell metabolic gene signatures across tumor-regions. Analysis performed on singlet T-cells based on HTO expression. Points represent individual samples from N=5 patients, with lines connecting paired samples from the same patient across regions. Pairwise comparisons were conducted using t-test (paired) or Wilcoxon test (paired) as appropriate*. Adjusted p-values for pairwise comparisons are indicated. **(H)** Heatmap of normalized cluster fractions across tumor-regions. Clustering used Euclidean distance on rows (cell clusters) and columns (tumor-regions). T-/ NK-cell cluster fractions are of total T-/NK-cells atlas, and myeloid/B cell fractions as of total myeloid/ B cells atlas. Analysis performed on singlets based on the HTO expression. Data include n=5 patients/samples per tumor-region. **(I)** Spearman correlation heatmap of immune cluster abundances. Positive correlations in red, negative in blue, with white indicating no correlation. Hierarchical clustering performed on both rows and columns using 1 – Spearman correlation and average linkage. * Statistical tests were chosen based on the normality of the data, with non-parametric tests applied for non-normally distributed data and parametric tests for normally distributed data (as assessed by Shapiro-Wilk).

Tissue-residency T cell markers were enriched in Tpex (CD8_Trm_CCL4) and CD8_Tex/Trm_ZNF683 cells, whereas CD8_Tex/Tcirc_GZMK cells overexpressed circulating T cell gene signatures (*20*) (Figure 4B and Supplemental Table 3 and 5). These findings are in line with recent studies showing two distinct differentiation pathways toward CD8 T cell exhaustion, of resident or circulating origin (*20, 21*).

CD4 T cells included naïve/memory (CD4_Tn_SELL, CD4_Tm_KLF2), Th17-like (CD4_Th_CD40LG), Tfh (CD4_Tfh_CXCL13), and Tregs (CD4_Tregs_FOXP3), with distinct gene expression profiles (Figure 4C). Four NK clusters were identified also, reflecting NK1 (NK_FCGR3A; e.g., *FCGR3A*, *FGFBP2*, *GZMB*, *SPON2*, *CCL3*, *CCL4*, *KLRD1*), NK2 (NK_NKG7, NK_TRDC; e.g., *XCL1*, *XCL2* and *AREG*), and NK3 (NK_TYROBP; e.g., *PPDPF*, *S100A6* and *IL32*) states (*38*) (Supplemental Figure 4E and Supplemental Table 5). NK cells, co-cluster partially with a small subset of effector-like T cells, due to co-expression of effector genes (e.g., *NKG7*, *FCGR3A*) (Supplemental Figure 4F). Cycling (CD4/CD8_Cycling_MKI67) and interferon-stimulated (CD4/CD8_IFN_ISG15) clusters were also identified, primarily derived from Tregs, Tex, and NK cells or Th cells respectively (Supplemental Figure 4, G and H).

Spatial analysis of T cell states revealed enrichment of naïve and memory CD4 T cells at the invasive margin (Figure 4D and Supplemental Figure 4, I and J), with a trend for Tregs and cycling T cells, in the tumor core (fig. S4, I and J). Independently of clustering, core T cells upregulated T cell exhaustion (e.g. *PDCD1*, *TIGIT*, *ID2*, *ENTPD1*, *HAVCR2*) and tissue residency (e.g. *ITGAE*, *ZNF683*) markers (Figure 4E). Expression of *KLRG1*, a marker of terminal-effector-memory (Temra) and circulating T cell states (*39, 40*), was increased in intermediate regions (Fig. 4E), whereas peripheral T cells expressed naïve/ memory genes (e.g. *KLF2*, *TCF7*, *IL7R*, *SELL*, *CCR7*, *LEF1*) (Figure 4E). Core CD8 T cells displayed tissue-resident and terminally exhausted signatures (CD8_Terminal_Tex), while peripheral CD8 T cells exhibited naïve/ early differentiated programs (Figure 4F, upper panel and Supplemental Table 3). Consistently, peripheral CD4 T cells were enriched in naïve and memory signatures, while core CD4 T cells expressed activated Treg signatures (CD4_TNFRSF9^+^_Treg) (Figure 4F, bottom panel and Supplemental Table 3).

These results indicate that TNFRSF9^+^ Tregs and Tex CD8 T cells tend to accumulate in central tumor regions (Figure 4F). Both signatures were previously linked to tumor reactivity (*21*). Supporting this, cycling cells were predominantly Tregs and exhausted CD8 T cells in the tumor core (Supplemental Figure 4K). These results suggest an enrichment of cycling, tumor-reactive T cells with exhaustion features in the tumor core, contrasted by naïve and memory-like T cell populations at the periphery.

To further investigate T cell functional differences across tumor regions, we analyzed the expression of metabolic signatures, including fatty acid (FA) metabolism, oxidative phosphorylation (OXPHOS), and TGF-β signaling pathways (Figure 4G and Supplemental Table 3). These programs were upregulated in T cells within the tumor core (Figure 4G), suggesting metabolic reprogramming under immunosuppressive conditions. Together, these data reveal regional distinct T cell states and metabolic adaptations, with a core tumor region dominated by exhausted, but tumor-reactive, proliferative and metabolically adapted T cells, emphasizing the functional and spatial organization of antitumor T cell responses within the tumor.

### Interplay of immune cell states across tumor regions

To understand how these spatially distinct T cell states interface with other components of the TME, we next examined correlations and co-localization patterns between T cell, NK cell, B cell and myeloid cell populations across tumor regions. Unsupervised hierarchical clustering of cell abundancies revealed distinct immune profiles across tumor regions, confirming pronounced spatial heterogeneity (Figure 4H). Central regions were dominated by M2-like and MMP-related macrophages, NK cells, Tregs, Tfh CD4 T cells, and circulating, Tex CD8 T cells (CD8_Tex/Tcirc_GZMK), consistent with an immunosuppressive environment (Figure 4H). In contrast, areas near the invasive margin, contained B cells, monocytes, DCs, M1-like macrophages, and naïve/memory CD4 T cells, suggesting a more active immune surveillance network.

Spearman correlation analysis of cell abundances revealed positive correlations among Tregs, Tfh CD4 T cells, cycling T cells, and circulating, Tex CD8 T cells (Figure 4I), populations enriched in central regions (Figure 4H). Conversely, DCs correlated with naïve and memory CD4 T cells, tissue-resident Tpex CD8 T cells and monocytes (Figure 4I), populations enriched in the periphery (Figure 4H). These results highlight coordinated immune programs within spatially defined niches, indicating a tumor core populated by immunosuppressive and tumor-reactive cells, while the periphery supports immune surveillance via APCs and memory T cells.

### Distinct TCR clonal expansion patterns across tumor regions

Given that Tex CD8 T cells and TNFRSF9^+^ Tregs cycle and bear tumor-reactive TCRs (*21*) and that both populations in our study tend to accumulate in the tumor core (Figure 4F), we reasoned that T cells in the tumor core should exhibit increased clonal expansion compared to peripheral T cells. To test this, we integrated TCR sequencing data with scRNA-seq data, defining T-cell clonotypes based on the CDR3β sequence. Our analysis revealed that Treg, Tfh, Trm and Tex populations contain the most expanded TCR clones (Figure 5A). Consistent with their tissue-resident versus circulating origins, TCR sharing was higher between CD8_Tex/Trm_ZNF683 and CD8_Trm_CCL4 states, than with CD8_Tex/Tcirc_GZMK (Supplemental Figure 5A).

**Figure 5.**
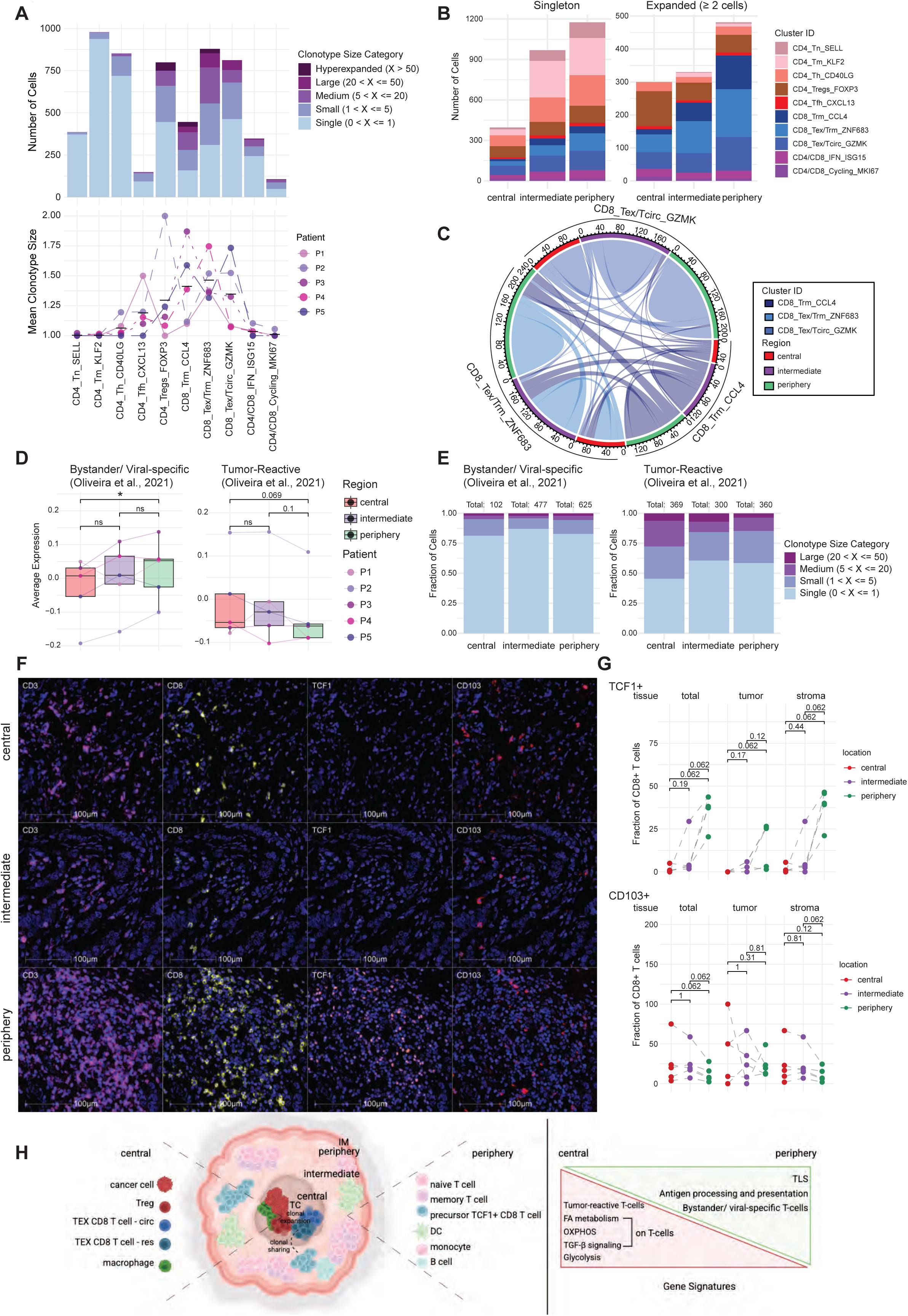
TCR clonal dynamics across tumor-regions in luminal breast cancer. **(A)** (upper panel): Clonal distribution across T-cell clusters. Clonotypes categorized by total cell count into singleton, small (2–5 cells), medium (6–20 cells), large (21–50 cells), and hyperexpanded (over 50 cells). A clonotype can appear in multiple clusters if its cells are distributed across them; color indicates expansion status across the entire dataset. (bottom panel): Mean clonotype size per patient in each cluster. Each point represents the mean clonotype size for a single patient within a given cluster. Analysis performed on total T-cells. **(B)** Distribution of singleton (one cell) and expanded (≥ 2 cells) clonotypes across T-cell clusters per tumor-region. Colors indicate relative abundance of T-cell clusters. Analysis performed on singlet T-cells based on HTO expression. **(C)** Circos plot of clonal sharing among CD8 T-cell clusters and tumor-regions, calculated by Jaccard index. Analysis performed on singlet CD8 T-cells based on HTO expression. **(D)** Expression of bystander/ viral-specific (left) and tumor-reactive (right) T-cell signatures across tumor-regions. Analysis performed on singlet T-cells based on HTO expression. Points represent individual samples from N=5 patients; lines connect paired samples. Pairwise comparisons were conducted using t-test (paired) for the bystander/ viral-specific signature expression analysis and the Wilcoxon (paired) test for the tumor-reactive signature expression analysis. Statistical tests were chosen based on the normality of the data, with non-parametric tests applied for non-normally distributed data and parametric tests for normally distributed data (as assessed by Shapiro-Wilk). Adjusted p-values are indicated; p ≤ 0.05 (*); ns= not significant. **(E)** Clonotype size distribution of bystander (identified as T-cells showing expression of bystander signature >1 & tumor-reactive signature <1) and tumor-reactive T-cells (identified as T-cells showing expression of bystander signature <1 & tumor-reactive signature >1) across tumor-regions. Total cell numbers per region indicated. A clonotype can appear in multiple clusters if its cells are distributed across them; color indicates expansion status across the entire dataset. **(F)** Representative IHC images of CD3 (pink), CD8 (yellow), TCF1 (orange) and CD103 (red) cells across regions. Scale bar: 100um. **(G)** Fraction of TCF1⁺ CD8⁺ (upper) and CD103⁺ CD8⁺ (bottom) T cells out of total CD8⁺ T cells across regions. Individual patients represented by dots; lines connect matched samples. Statistical comparisons between tumor-regions were conducted using paired Wilcoxon test; p-values are indicated. **(H)** Graphical model summarizing spatial T-cell dynamics. Created with BioRender.com.

Analysis of TCR clonal expansion patterns across tumor regions revealed a trend—though not statistically significant—toward higher expansion in the tumor core, as indicated by a greater fraction of large clones (Supplemental Figure 5B), larger mean clonotype size (Supplemental Figure 5C), and lower Shannon diversity index (Supplemental Figure 5D), relative to the periphery.

To further dissect these patterns, we stratified clones into singletons (detected in one cell) and expanded (detected in ≥ 2 cells) clones. We observed that singletons were enriched in naïve and memory CD4 T cells, while expanded clones were mainly Tregs and CD8 T cells (Figure 5B and Supplemental Figure 5E). Notably, expanded Treg clones were predominantly in the tumor core (Figure 5B and Supplemental Figure 5E). In contrast, we observe a reduction in clonally expanded Tpex CD8 T cells (CD8_Trm_CCL4) toward central regions, with expanded CD8 clones in the core deriving mainly from Tex states (Figure 5B). These findings indicate region-specific clonal dynamics: the tumor core harbors higher fractions of clonally expanded Treg and Tex T cells, whereas the periphery of the tumor is enriched in diverse, less-expanded naïve and memory T cells.

Clonal sharing analysis further revealed lineage relationships between peripheral and central CD8 T cell populations. Tissue-resident Tpex CD8 T cells (CD8_Trm_CCL4) in the periphery shared clones with tissue-resident Tex CD8 T cells (CD8_Tex/Trm_ZNF683) in the core, suggesting a local differentiation trajectory from peripheral precursors to terminally exhausted resident CD8 T cells in the tumor core (Figure 5C). In contrast, circulating Tex CD8 T cells (CD8_Tex/Tcirc_GZMK) showed clonal overlap across regions but minimal sharing with resident populations (Figure 5C). These findings suggest that while tissue-resident exhausted CD8 T cells arise locally, circulating exhausted CD8 T cells constitute a distinct lineage, with limited interconversion between circulating and resident compartments.

The results presented so far suggest that tumor-reactive T cells are enriched and exhibit greater clonal expansion in the tumor core. To test this, we analyzed tumor-reactive versus bystander T cell gene signatures (*41*) across tumor regions (Supplemental Table 3). Bystander signatures were enriched in peripheral T cells, while tumor-reactive signatures were elevated in the core (Figure 5D).

Next, we stratified T cells based on these signatures and examined their clonal expansion and spatial distribution. Analysis of bystander versus tumor-reactive T cell abundances across regions showed dominance of bystander T cells in the periphery, whereas tumor-reactive T cells peaked in the core (Supplemental Figure 5F). Clonal expansion analysis further revealed that bystander T cells remained largely non-expanded or oligoclonal across all regions, whereas tumor-reactive T cells exhibited greater clonal expansion, with the highest fraction of large clones observed in the tumor core (Figure 5E). These findings support the hypothesis that the tumor core harbors a concentrated population of clonally expanded, tumor-reactive T cells, whereas the periphery is enriched in non-expanded bystander populations. Public viral-specific TCR clones recognizing influenza epitopes were detected only in tissue-resident CD8 T cells from peripheral or intermediate, but not central tumor regions (Supplemental Figure 5G).

Finally, multiplex IHC validated these spatial T cell patterns. TCF1⁺ CD8 T cells, marking Tpex/ memory-like cells, were significantly enriched in peripheral regions (Figure 5, F and G), while CD103⁺ CD8 T cells were detected across all tumor regions, indicating the presence of tissue-resident memory T cells throughout the TME (Figure 5, F and G). Collectively, these findings define a spatial T cell architecture with a precursor-, memory-rich periphery and a clonally expanded, tumor-reactive core.

## DISCUSSION

Our results reveal a spatially organized immune landscape in luminal breast cancer. Unlike conventional spatial transcriptomic technologies, which preserve tissue architecture, but lack full single-cell resolution or TCR information, our approach allowed us to simultaneously capture the functional transcriptome and T cell clonal architecture across discrete tumor regions. Here, by profiling three ex-vivo biopsies from central, intermediate and peripheral regions with single-cell 5’ V(D)J and gene expression analysis coupled to protein hashtag barcoding, we retained coarse spatial context while enabling high-resolution mapping of cell states and clonal dynamics. This approach uncovered patterns of immune suppression and activation shaped by local tumor microenvironmental cues.

We demonstrate spatial compartmentalization of myeloid and T cell states and TCR clonality across tumor regions, suggesting a working model in which immunosuppression is established from the periphery toward the core, through progressive enrichment in M2-like macrophages and Tregs, coupled with spatially restricted T cell activation and exhaustion (Figure 5H). What, then, drives these changes in spatial environments within the tumor?

This spatial compartmentalization likely reflects both differential immune infiltration and local cues from the TME. Central regions, enriched in cancer cells, are likely more hypoxic and metabolically reprogrammed (*42*), with elevated suppressive cytokines, such as TGF-β and IL-10 (*43–45*), immunomodulatory metabolites, like adenosine and lactate (*46–48*), steroid (e.g., estrogens, androgens, cortisol) (*49–52*) and growth factors (*53, 54*). Progressive monocyte differentiation into M2-like macrophages, along with chronic TCR engagement on Tregs and Tex may reinforce immunosuppression through self-sustaining feedback loops.

The novelty of our study lies in spatially mapping these transitions. Naïve T cells, monocytes, DCs and B cells are enriched near the invasive margins, indicative of recent tumor infiltration, likely via vascular structures, such as endothelial cells and pericytes (*55*), and are linked with antigen presentation and the activation of T cells. This aligns with upregulated TLS and APC signatures in these regions, both of which are known to support T cell priming and activation (*30*). The spatial distribution of these immune cells, along with the presence of TLS, suggests that the periphery of the tumor may harbor an active immune surveillance network, contributing to the initiation of immune responses.

Conversely, the tumor core is dominated by M2-like macrophages, Tregs and Tex CD8 T cells. T cells in these regions upregulated gene programs related to FA metabolism and OXPHOS, as well as TGF-β signaling, consistent with metabolic adaptation to a nutrient- and oxygen-poor, immunosuppressive microenvironment (*56–58*).

We show that T cells in the tumor core may recognize tumor antigens. Tumor-reactive CD8 and CD4 T cell gene signatures (*21*) were upregulated in the core, consistent with increased clonality, a proxy of tumor reactivity (*59, 60*), in Tregs and Tex cells in central regions. In line with these observations, peripheral T cells displayed bystander, viral-specific characteristics. In regions near the invasive margin, T cells are exposed to a less suppressive environment, encounter fresh tumor antigens presented by DCs or within TLSs, and retain stem-like properties, with increased TCF1 expression, indicative of a memory-like phenotype (*18, 61*), as validated by multiplex IHC analysis.

Together, these findings indicate a spatial organization of T cell states and TCR specificities: the tumor periphery is immunologically active, populated by precursor T cells and APCs, while the core is an immunosuppressive niche harboring clonally expanded, tumor-reactive T cells with exhaustion features. These results indicate a complex interplay between immune suppression and activation within the TME, suggesting that targeting both tumor-specific, exhausted T cells in the core, and stem-like, immune-active, components at the margins could enhance anti-tumor immunity.

While we made efforts to capture the intratumor spatial immune heterogeneity, our study has limitations. Transcriptomic and TCR sequencing do not fully account for the complexities of protein expression, cell-cell interactions, or cytokine profiles, which could further refine understanding of immune dynamics. Furthermore, although our findings suggest consistent immune dynamics across tumor regions, they are based on a single tumor type and may not generalize across cancers. Future studies integrating multi-omics, in vivo models, and therapeutic interventions are needed to validate these observations and explore therapeutic potential the of targeted modulation of both exhausted, tumor-reactive and immune-active T cell populations.

Ultimately, this work provides a high-resolution view of spatial immune dynamics within luminal tumors, highlighting the balance between core immunosuppression and peripheral immune activation. Integrating immune cell states with clonal dynamics, we propose a refined model for spatial immune remodeling, offering a foundation for further exploration of the tumor immune landscape. Future research integrating protein-protein interactions, immune factors, and cytokine profiles within the TME will expand upon these insights, offering new avenues for therapeutic strategies. This work could pave the way for the development of more targeted treatments to reprogram the tumor microenvironment to enhance anti-tumor immunity.

## METHODS

### Human samples and Tumor collection

Tumor tissue samples were collected ex vivo from five female patients with early-stage luminal breast cancer, immediately following surgical resection. For each tumor, three macroscopically distinct regions were sampled: one proximal to the tumor core, one near the invasive margins, and one intermediate between the two. The study was conducted in compliance with the Declaration of Helsinki and the national regulatory requirements. Study eligible patients were over the age of 18, had been diagnosed with early-stage luminal breast cancer and had not been previously treated. Patient demographics and tumor characteristics are summarized in Supplementary Table S1.

### Single-cell suspension and labeling

Tissues were enzymatically and mechanically dissociated into single-cell suspensions, followed by red blood cell and dead cell removal. Cells were subsequently labeled by oligonucleotide-barcoded antibodies (HTO) to enable samples multiplexing. Viability and cell counts were measured before loading on the 10X Genomics Chromium platform. Detailed dissociation and HTO protocols are provided in Supplementary Materials and Methods.

### Single-cell RNA-, CITE- and TCR-sequencing profiling

Single-cell suspensions were processed using the Chromium Single Cell 5’ Reagent Kit with Feature Barcoding. Libraries were prepared and sequenced according to manufacturer protocols. Library preparation specific, quality control, and sequencing parameters are included in Supplementary Materials and Methods.

### Data processing and analysis

Raw sequencing data were processed using Cell Ranger (v6.0.2) for demultiplexing, alignment to human genome reference GRCh38 and gene counting. Downstream analyses were performed using *Seurat* (*62, 63*) (v4.2.1; v4.3.0; v5.0.1) for normalization, clustering and dimensionality reduction, and *harmony* (*64*) for batch correction. Detailed description of cell type annotation, gene signature scoring, pseudobulk analysis and label transfer provided in Supplementary Materials and Methods.

### TCR sequencing and clonal analysis

TCR sequences were aligned, annotated, and clonotypes defined based on CDR3β amino acid sequences. Clonal expansion, diversity, and spatial distribution analyses were performed as described in the main text. Detailed methods for clonal analysis provided in Supplementary Materials and Methods.

### Multiplex Immunohistochemistry (mIHC)

Formalin-fixed, paraffin-embedded matched tumor blocks were stained with multiplex panels targeting immune populations. Imaging, spectral unmixing, and quantitative analysis were performed using Vectra 3 and HALO software. Complete antibody panels and staining workflows are provided in Supplementary Materials and Methods.

### Statistical analysis

All statistical analyses were performed using R v4.2.2. For the single-cell analysis samples were treated as pseudobulks using the AggregateExpression() function of Seurat package. The specific test used is indicated in the figure legend and statistically significant *p* values are shown in graphs (a *p* < 0.05 was considered as statistically significant). For cluster abundance comparisons across tumor regions, two-way ANOVA with Geisser-Greenhouse correction was performed, followed by Tukey’s post-hoc test for pairwise comparisons. For comparisons of discrete variables between groups (e.g., tumor core vs. peripheral regions), normality was assessed using the Shapiro-Wilk test and then Wilcoxon rank-sum tests were applied for non-parametric data, while t-tests were used for normally distributed data. For statistical evaluations the following R packages were utilized: afex_1.4-1; emmeans_1.11.0; stats_4.2.2; ggpubr_0.6.0; ggsignif_0.6.4.

## Supporting information

Supplemental Data (inc. Figures & Methods)

## Data availability

The data for this study are deposited in the Gene Expression Omnibus database (GSE260509). Data are currently private, will be shared with the reviewers upon request, and will be publicly available upon publication.

## LIST OF SUPPLEMENTARY MATERIALS

Supplementary Materials and Methods.

Supplementary Figures 1-5.

Supplementary Tables 1-5.

## AUTHOR CONTRIBUTIONS

SA conceived and supervised the project. CM designed and performed experiments, analyzed and interpreted data. PEB and VK assisted with data analysis. LL and AN assisted with multiplex IHC. ER, LD and AVS assisted with provision of clinical samples. PS supported clinical samples selection. OL and ER contributed analytical input. CM and SA wrote the original manuscript. All authors reviewed and edited the final manuscript.

## ACKNOWLEDGMENTS

High-throughput sequencing was performed by the ICGex NGS platform of the Institute Curie and by VSC (Flemish SuperComputer Center). The computational resources and services for demultiplexing and aligning the samples were performed by the Bioinformatics platform of the Institute Curie and by VSC. The authors would like to thank the Institute Curie ICGex NGS platform for high-throughput sequencing. The authors thank Sylvain Baulande, Sonia Lameiras, Patricia Legoix, and Benoit Albaud from ICGEX for sample preparation, data acquisition, and technical advice. The authors thank Josh J. Waterfall for critically reading the manuscript. The authors would like to thank the pathologists and surgeons from the Department of Pathology of Institute Curie for patient sample collection and coordination, as well as all patients and their families.

