## Supplemental Data (inc. Figures & Methods) for "Analysis of intratumoral immune heterogeneity reveals spatial organisation of immunosuppression in breast cancer"

**The PDF file includes:**
Materials and Methods.
Supplemental Figures 1 to 5.
Description for Supplemental Tables 1 to 5.

#### Materials and Methods

##### Human samples

Tumor tissue samples were collected ex vivo from five patients with early-stage luminal breast cancer, immediately following surgical resection. All study participants are females. The study was conducted in compliance with the Declaration of Helsinki and the national regulatory requirements. Study eligible patients were over the age of 18, had been diagnosed with early-stage luminal breast cancer and had not been previously treated. Demographic and tumor characteristics of the study population were obtained and are included in Supplementary Table 1.

##### Ex-vivo tumor sampling

For each patient, three macroscopically distinct regions of the tumor were identified: one proximal to the tumor core, one near the invasive margins, and one intermediate between the two. Fresh tissue from each region was excised and immediately processed for further analysis.

##### Biopsy tissue dissociation

Tumor-tissue samples were cut in about 0.5 mm3 pieces using scalper. Tissues were digested enzymatically by a 20’ incubation, at 37°C under agitation, in CO2-independent medium (Invitrogen) with collagenase I (2 mg/ml; #637958; Sigma-Aldrich), hyaluronidase (2 mg/ml; #H3506; Sigma-Aldrich), and deoxyribonuclease (25 ug/ml; #D5025; Sigma-Aldrich). The tissue pieces were triturated with a 20mL syringe plunger on a 70um cell strainer (BD), washed with 1X PBS (Invitrogen) with 1% fetal bovine serum (FBS) and 2mM EDTA (Gibco), until uniform cell suspensions were obtained. Suspended cells were subsequently centrifuged for 10’ at 300 rcf at 4°C.

##### HTO labeling

Antibody-mix supernatant preparation was performed by adding 1ul of the HTO antibody in 50ul of 1X PBS (Invitrogen) with 1% bovine serum albumin (BSA). The mix was subsequently centrifuged at 14,000 rcf. Supernatant was transferred to a new tube and kept at 4°C.

Cells were resuspended in 1mL of 1X PBS (Invitrogen) with 0.04% (BSA) and centrifuged at 150rcf for 10’ at 4°C. Supernatant was removed and cell pellet was resuspended in 50ul chilled 1X PBS (Invitrogen) with 0.04% BSA. 5ul of Human TruStain FcX™ (Fc Receptor Blocking Solution) was added on top and cells were incubated for 10’ at 4°C. 50ul of the prepared antibody-mix supernatant was added on top and mix was gently pipetted 10x (pipette set to 90ul). Cells were subsequently incubated for 30’ at 4°C without light exposure. Next, cells were washed 3x with 1mL chilled 1X PBS (Invitrogen) with 0.04% BSA followed by a centrifuge at 150 rcf for 10’ at 4°C. Upon washing, cells were counted, and proper volumes were obtained per sample for mixing of equal number of cells per HTO-labelled sample.

**Red blood cell - Dead cell removal and single cell preparation**

Upon HTO labeling, red blood cells and dead cells were removed using magnetic isolation kit by StemCell technologies, following manufacturer instructions (product numbers: #18170RF and #17899 respectively). Upon purification, cells were resuspended in 1X PBS with 0.04% BSA, in a concentration of 1,000 cells/ ul for a targeted cell recovery of 10,000 cells as indicated by 10X Genomics. Cell numbers and viability were measured using a Countess II Automated Cell Counter (Thermo Fisher Scientific) and hemocytometer/ trypan blue.

**Single-cell RNA-, CITE- and TCR- sequencing profiling**

Single-cell suspensions were loaded onto a Chromium Single Cell Chip (10X Genomics) according to the manufacturer’s instructions for co-encapsulation with barcoded gel beads at a target capture rate of 10,000 individual cells per sample, based on the initial number of cells. For all samples RNA, protein and TCR libraries were synthetized by following the Chromium Single Cell 5' Reagent Kits User Guide (v2 Chemistry Dual Index) with Feature Barcoding technology for Cell Surface Protein and Immune Receptor Mapping. cDNA for both V(D)J/ 5’ gene expression library and cell surface protein/ immune receptor mapping library were profiled using both Qubit (ThermoFisher scientific) and Bioanalyzer High Sensitivity DNA kit (Agilent Technologies). Libraries for RNA-seq, cell surface protein and V(D)J were prepared following the manufacturer’s user guide (10X Genomics), then profiled using Kapa Library Quantification kit (Kapa Biosystems) and quantified with Qubit (ThermoFisher Scientific) and Tapestation (Agilent Technologies). Libraries were sequenced with Novaseq (Illumina). All sequencing was done according to the manufacturer’s specification (10X Genomics).

**Single cell sequencing data analysis**

Single-cell expression was analyzed using the Cell Ranger Single Cell Software Suite (10X Genomics, v6.0.2) to perform quality control, sample demultiplexing, barcode processing, and single-cell 5′ gene counting. Sequencing reads were aligned to the GRCh38 human reference genome (GRCh38-2020-A) and quantified using cellranger count function. Filtered gene barcodes matrices, containing barcodes with UMI counts passing threshold for cell detection, were used for further analysis. Empty droplets removal was performed by the EmptyDrops package *(1)*, implemented in CellRanger. To mitigate potential batch effects, while maintaining biological differences across samples, we performed data integration using harmony *(2)*.

Downstream analysis was performed using Seurat *(3)* (version 4.2.1; 4.3.0; 5.0.1)· Quality control was performed to exclude cells with less than 200 or more than 6000 detected genes and with greater than 15% mitochondrial RNA content (for patient samples P1 and P2) or greater than 10% mitochondrial RNA content (for patient samples P3-P5). Filtered cell barcodes from CellRanger output were used as whitelist to quantify HTOs using CITE-seq-Count (v1.4.5).

For cell clustering, raw UMI counts were log normalized, and variable genes called on each dataset independently. The top 6000 variable features were identified using the "vst" method from Seurat. Variable T cell receptor and immunoglobulin genes were removed from the list of variable genes to prevent clustering based on variable V(D)J transcripts. We used Harmony *(2)* to correct batch effects across patient-samples. Cell cycle score for S and G2/M cell cycle phase was assigned to each cell, based on previously published datasets *(4)* using the CellCycleScoring function.

Scaled scores for each gene were calculated using the Scale Data function and regressed against number of UMIs per cell and mitochondrial RNA content. Scaled data was used an input into PCA based on variable genes. A stressed-induced cluster of low-quality cells (highly expressing mitochondrial/ MT- and heat-shock-protein/HSP- genes) was removed from the initial analysis and cells were re-analyzed. Clusters were identified based on the shared nearest neighbor (SNN) clustering on the first 20 PCs and resolution= 0.5. Principal components were used to generate UMAP projections· Unique cluster-specific differential expressed genes were identified by performing the Seurat FindAllMarkers function using Wilcoxon test on the RNA assay. Signature scores were computed using the Seurat function AddModuleScore using the gene signature of interest and setting the number of control genes from the same bin of expression at 5. This function calculates for each individual cell the average expression of each gene signature, subtracted by the aggregated expression of control gene sets matched for individual gene expression level.

**Pseudobulk analysis**

To correct for the sparsity and heterogeneity of single-cell data, we performed pseudobulk analysis. We used the the 'AggregateExpression' function of Seurat, that returns summed counts ("pseudobulks") for each identity class. Next, mean gene signature expression per sample was plotted.

**Label transfer using a reference**

Label transfer was performed using Seurat v3 FindTransferAnchors and TransferData functions. FindTransferAnchors function identifies a set of anchors between a reference and query object and next TransferData function uses the set of anchors computed to transfer categorical or continuous data from the reference to the query object. Cycling cells and interferon-induced T cells cluster together due to co-expression of cell cycle genes and interferon-stimulated genes (ISGs) respectively. To identify their identity, label transfer method was used to assign the cells from the cycling cluster and IFN cluster into their cluster-of-origin on the broad cluster UMAP and on the T cell cluster UMAP.

**Single cell TCR-sequencing data processing**

TCR reads were aligned to the GRCh38 reference genome and consensus TCR annotation was performed using cellranger vdj (vdj_GRCh38_alts_ensembl- version 3.1.0-3.1.0). TCR annotation was performed using the 10X cellranger vdj pipeline. Doublets were excluded based on >1 β chain detected. Clonotypes assignment and downstream analysis of the clonotype dynamics was performed by scRepertoire package *(5)* (version 1.7.2) and custom code. Clonotypes were defined based on the shared CDR3β amino acid sequence.

As a metric of T cell clonal expansion, we used mean clonotype size, which was computed as the ratio between the total number of cells and the total number of unique clonotypes. To assess TCR diversity across tumor regions, we calculated the Shannon diversity index, which accounts for both clonal richness and evenness. The index was normalized by sample size to correct for differences in sequencing depth. In addition, we quantified clonal expansion by computing the mean clone size per region, normalizing it by the total number of T cells in each sample. Statistical comparisons between tumor regions were performed using the Kruskal-Wallis test for global differences and Wilcoxon paired tests for pairwise comparisons.

In Fig.5A, 5E and Supplementary Fig. 5B the distribution of each clonotype across tumor regions and clusters was visualized, with clonotypes categorized into expansion groups based on their total abundance across the entire dataset. Each clonotype was assigned to a group corresponding to its expansion status (e.g., small, medium, large etc). Categorizing clonotypes based on their total abundance across the entire dataset allows a consistent classification of expansion status and a clear representation of how clonotypes distribute across regions and clusters while maintaining their overall expansion category.

To identify viral-specific TCRs, we searched in VDJdb database *(6)* for exact matches (Levenstein distance = 0) between CDR3β sequences previously experimentally confirmed to recognize viral epitopes and the ones from our data. We focused on TCRs specific to epitopes from Influenza virus assuming that due to the virus' abundance some of the patients might have been exposed to it. We found 4 sequences with exact match for Influenza-specific TCRs in patients P3 and P4, all of them recognizing HLA-A*02:01-M158–66 epitope.

**Bystander vs. tumor-reactive T cells**

To analyze the spatial distribution and clonal expansion patterns of bystander vs. tumor-reactive T cells across regions in Fig. 5D-5E, we subsetted cells based on: expression of bystander/ viral-specific signature >1 & tumor-reactive signature < 1 was used to identify bystander/ viral-specific T cells and expression of tumor-reactive signature > 1 & bystander viral-specific signature <1 was used to identify tumor-reactive T cells *(7)*.

**Multiplexed immunohistochemistry**

Paraffin-embedded tissue blocks were cut with a microtome into fine slivers of 3 microns. Immunostaining was processed in a Bond RX automated (Leica) with Opal™ 7-Color IHC Kits (Akoya Biosciences, NEL821001KT) according to the manufacturer's instructions. The multiplex panel consisted of the following antibodies: panel 1: CD8 (Agilent, #M7103), CD3 (Agilent, #GA503), CD4 (Cell Signaling, #48274), FoxP3 (Diagomics Master Diagnostica, #MAD-000536QD-7), CD163 (Leica, #NCLCD163), pan Cytokeratin (Agilent, #M3515); panel 2: CD103 (Abcam, #ab129202), CD8 (Agilent, #M7103), CD3 (Agilent, #GA503), PD-1 (Abcam, #ab137132), CD39 (Abcam, #ab223842), pan Cytokeratin (Agilent, #M3515); panel 3: CD103 (Abcam, #ab129202), CD8 (Agilent, #M7103), CD3 (Agilent, #GA503), TCF1 (Cell Signaling, #2203), Granzym K (Abcam, #ab282703), pan Cytokeratin (Agilent, #M3515).

Tissue sections were cover slipped with Prolong™ Diamond Antifade Mountant (ThermoFisher scientific) and stored at 4°C. Subsequently, slides were scanned using the Vectra ® 3 automated quantitative pathology imaging system (Vectra 3.0.5; Akoya Biosciences). Multispectral images were unmixed and analyzed using the inForm Advanced Image Analysis Software (inForm 2.6.0; Akoya Biosciences) and the HALO software for immune subsets quantification.

### Supplementary Figure 1

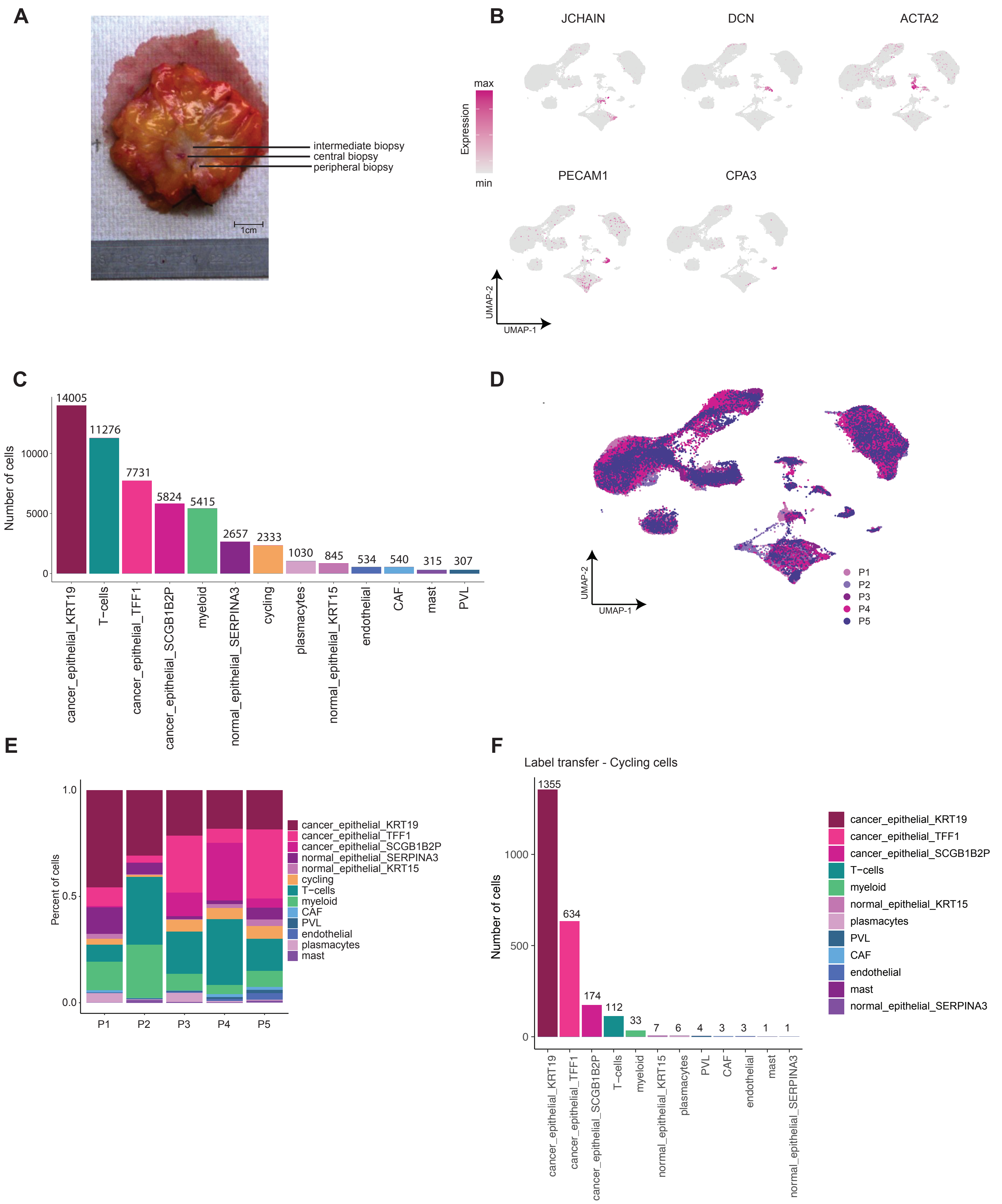

**Supplemental Figure 1. Sample preparation and detailed single-cell sequencing data information, related to Figure 1. (A)** Representative macroscopic image of sample retrieval. **(B)** Feature plot showing the normalized expression of marker genes in tumor-infiltrating cells. **(C)** Number of cells per cluster. **(D)** UMAP visualization of 52,812 tumor-infiltrating single cells, color-coded by patient. Each point represents a single cell, with colors distinguishing cells from different patients. **(E)** Relative abundance of cell clusters per patient. **(F)** Number of cycling cells across identified clusters. Cycling cells' identity was identified through label transfer from the cycling cluster to all other clusters.

### Supplementary Figure 2

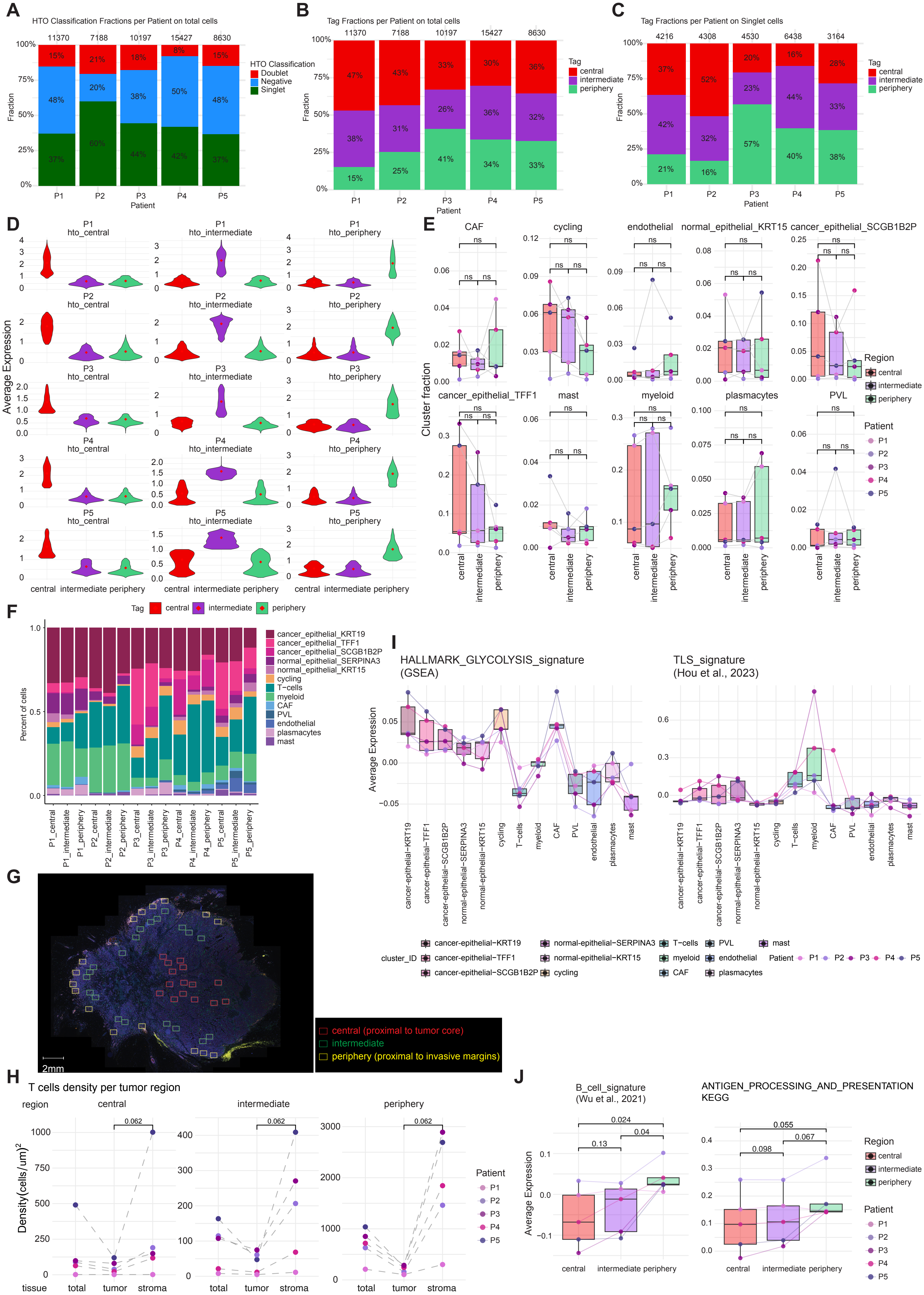

**Supplemental Figure 2. Detailed CITEseq data information, related to Figure 2. (A)** Bar plot showing relative cell abundances per patient, classified as singlets (labeled with a single hashtag), doublets (labeled with >1 hashtags), or negatives (low or undetectable hashtag expression) based on HTO expression. **(B)** Bar plot showing relative cell abundances per patient, assigned to central, intermediate or periphery tumor-region based on the hashtag-antibody expression on total cells. **(C)** Bar plot showing the relative abundance of cells per patient, assigned to central, intermediate or periphery tumor-region based on HTO expression on singlets. **(D)** Violin plot of average hashtag expression per sample and per patient on singlets. **(E)** Cluster frequencies per tumor region. Each dot represents a single sample (n = 15). Samples were derived from N = 5 patients, with 3 tumor-region samples (central, intermediate, and periphery) collected from each patient. Individual patients are represented by colored dots, and lines connect samples from the same patient. Statistical comparisons between tumor regions were conducted using a two-way repeated measures ANOVA with Greenhouse-Geisser correction. Tukey's post-hoc test was applied for pairwise comparisons. Adjusted p-values for pairwise comparisons are indicated; ns= not significant. Analysis performed on singlet cells based on hashtag-antibody expression. **(F)** Bar plot showing the relative abundance of cell clusters per sample (per patient and tumor-region). Analysis performed on singlet cells based on HTO expression. **(G)** Representative image of mIHC showing central (proximal to tumor core, red), intermediate (green), and peripheral (proximal to invasive margins, yellow) tumor regions selected for IHC analysis. Colored squares indicate areas selected for quantitate IHC analysis in each region. Selection was based on spatial location and morphological features to ensure representative sampling. Scale bar: 2mm. **(H)** CD3 T cell density (cells/ $um^2$) across different tumor regions: central- core, intermediate, and periphery-invasive margins and tissue: tumor tissue vs. stroma tissue. Data points represent individual patient samples, with each dot corresponding to a unique analysis region. Statistical comparisons were performed using the Wilcoxon signed-rank test; p-values are indicated. **(I)** Average expression of HALLMARK_GLYCOLYSIS (left panel) and TLS signature (right panel) across cell clusters. Points represent individual samples from N=5 patients, with lines connecting paired samples from the same patient across regions. Analysis performed on singlet cells based on HTO expression. **(J)** Expression of B cell signature (left panel) and Antigen_Processing_and_Presentation_KEGG signature (right panel) across tumor regions. Analysis performed on singlet cells based on HTO expression. Points represent individual samples from N=5 patients, with lines connecting paired samples from the same patient across regions. Pairwise comparisons were conducted using the t-test (paired).

#### Supplementary Figure 3

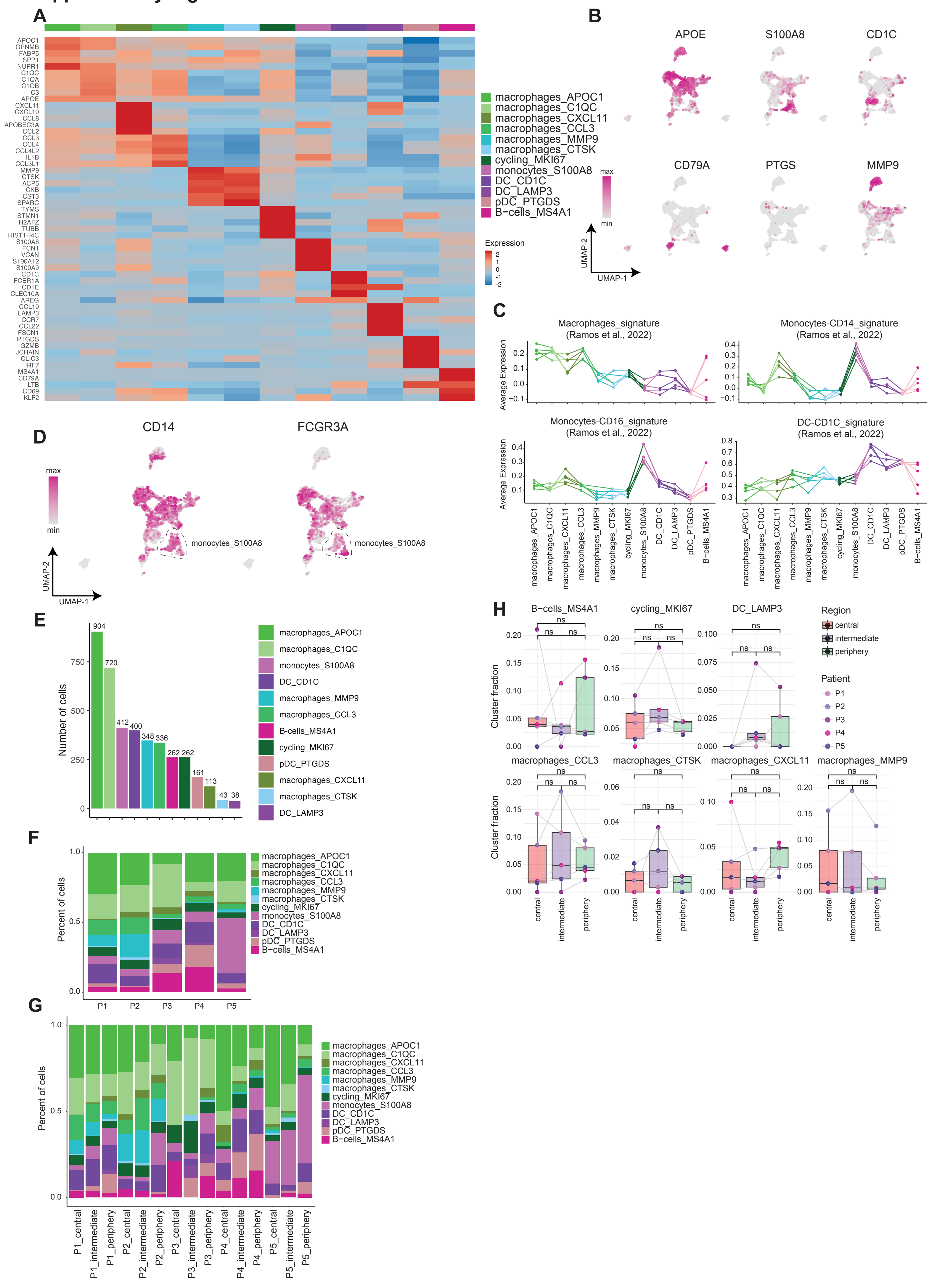

**Supplemental Figure 3. Detailed myeloid and B cell transcriptomic data, related to Figure 3. (A)** Heatmap of normalized expression of top5 differential expressed genes (DEGs) per myeloid cell cluster (wilcoxon rank sum test). **(B)** Feature plot showing the normalized expression of selected gene markers demonstrating myeloid cell type identity. **(C)** Average expression of myeloid-states gene signatures across myeloid clusters. Points represent individual samples from N=5 patients, with lines connecting paired samples from the same patient across regions. **(D)** Feature plot showing the normalized expression of selected gene markers demonstrating monocyte type identity: classical (CD14 expression) vs. non-classical (CD16/ FCGR3A expression). **(E)** Number of cells per myeloid cluster. **(F**) Bar plot showing the relative abundance of myeloid clusters per patient. **(G)** Bar plot showing the relative abundance of myeloid clusters per sample (per patient and tumor-region). Analysis performed on singlet myeloid cells based on hashtag-antibody expression. **(H)** Cluster frequencies per tumor region. Each dot represents a single sample (n = 15). Samples were derived from N = 5 patients, with 3 tumor-region samples (central, intermediate, and periphery) collected from each patient. Individual patients are represented by colored dots, and lines connect samples from the same patient. Statistical comparisons between tumor regions were conducted using a two-way repeated measures ANOVA with Greenhouse-Geisser correction. Tukey's post-hoc test was applied for pairwise comparisons; ns= not significant. Analysis performed on singlet myeloid cells based on hashtag-antibody expression.

#### Supplementary Figure 4

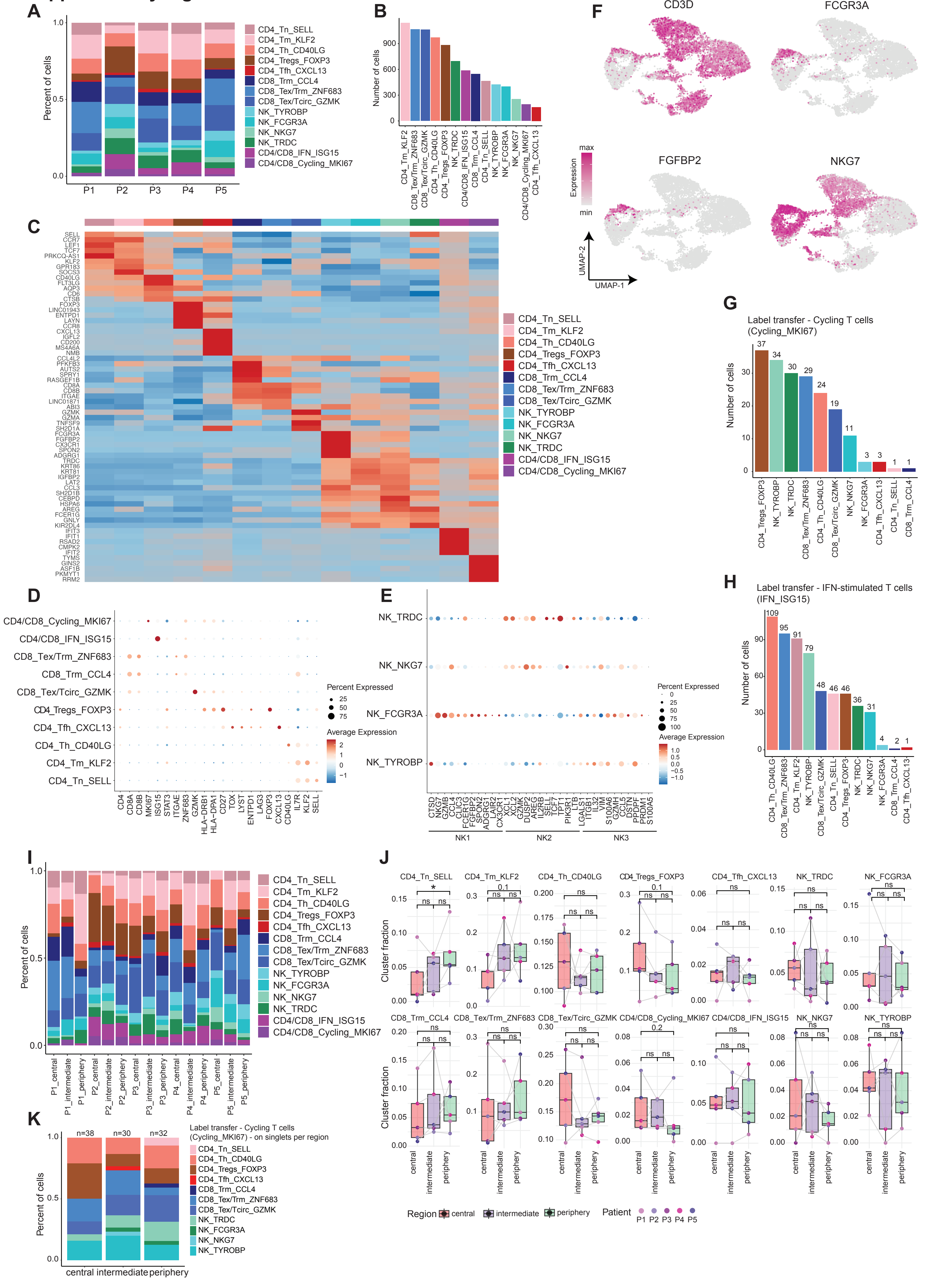

**Supplemental Figure 4. Detailed T- and NK- cell transcriptomic data, related to Figure 4. (A)** Bar plot showing the relative abundance of T-cell clusters per patient. **(B)** Number of cells per T-cell cluster. **(C)** Heatmap of normalized expression of top5 differential expressed genes (DEGs) per T-cell cluster (wilcoxon rank sum test). **(D)** Dot plot showing expression of marker genes demonstrating T-cell state across T-cell clusters. Dot size indicates the percentage of cells expressing the gene within the cluster. Color intensity represents the average expression level of the gene in the cluster. **(E)** Dot plot showing expression of marker genes demonstrating NK cell state across NK clusters. Dot size indicates the percentage of cells expressing the gene within the cluster. Color intensity represents the average expression level of the gene in the cluster. **(F)** Feature plot showing the normalized expression of selected genes. **(G)** Number of cycling T cells (CD4/CD8_Cycling_MKI67) across identified clusters. Cycling cells' identity was identified through label transfer of cells from the cycling cluster onto the rest of the T-cell clusters. **(H)** Number of interferon-stimulated T-cells (CD4/CD8_IFN_ISG15) across identified clusters. IFN cells' identity was identified through label transfer of cells from the IFN cluster onto the rest of the T-cell clusters. **(I)** Bar plot showing the relative abundance of T-cell clusters per sample (per patient and tumor-region). Analysis performed on singlet T-cells based on the hashtag-antibody expression. **(J)** T-cell cluster frequencies per tumor region. Each dot represents a single sample ($n = 15$). Samples were derived from $N = 5$ patients, with 3 tumor-region samples (central, intermediate, and periphery) collected from each patient. Individual patients are represented by colored dots, and lines connect samples from the same patient. Analysis performed on singlet T-cells based on the hashtag-antibody expression. Statistical comparisons between tumor regions were conducted using a two-way repeated measures ANOVA with Greenhouse-Geisser correction. Tukey's post-hoc test was applied for pairwise comparisons. Adjusted p-values for pairwise comparisons are indicated; $p \leq 0.05$ (*); ns= not significant. **(K)** Bar plot showing the relative abundance of cycling T-cells per tumor-region. Analysis performed on singlet T-cells based on hashtag-antibody expression.

#### Supplementary Figure 5

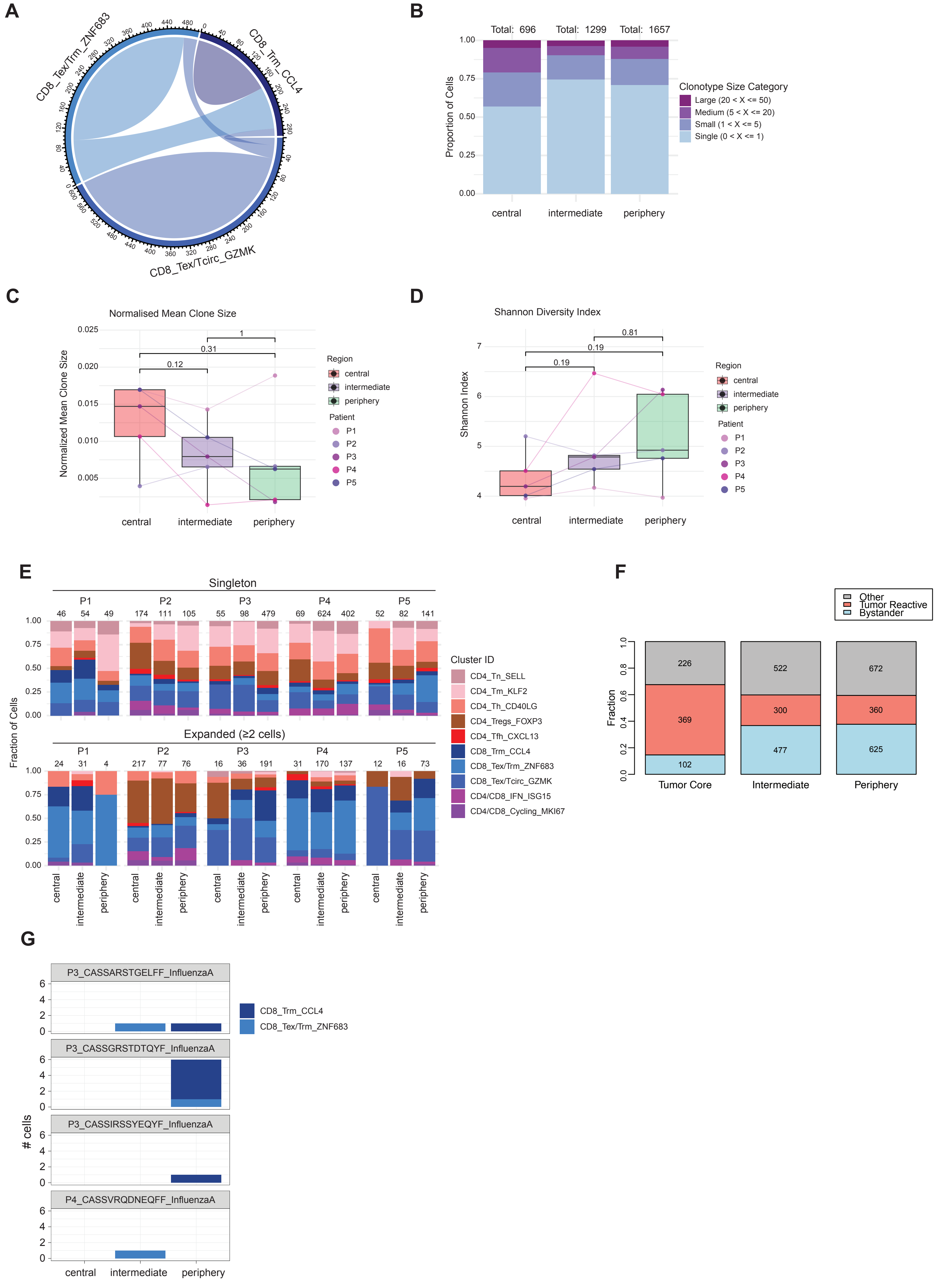

**Supplemental Figure 5. Spatial TCR profiling, related to Figure 5. (A)** Circos plot showing the clonal sharing among CD8 T-cell clusters. Clonal sharing was calculated based on the jaccard index. Analysis performed on total CD8 T-cells. **(B)** Bar plots showing the clonotype size distribution across tumor-regions. Total number of cells in each region is indicated in the top of each bar. The distribution of each clonotype's cells across tumor-regions was plotted, with clonotypes represented by different colors according to their size category. A clonotype can appear in multiple tumor-regions if its cells are distributed across them, and the color represents the clonotype's expansion status across the entire dataset. Analysis performed on singlet T-cells based on the hashtag-antibody expression. **(C)** Normalized mean clone size across tumor regions per patient. Mean clone size was normalized by the total cell size of each sample. Data points represent individual patient samples, with each dot corresponding to a unique analysis region. Analysis performed on singlet T-cells based on hashtag-antibody expression. Statistical comparisons were performed using the Wilcoxon test (paired); p-values are indicated. **(D)** Shannon diversity index across tumor regions per patient. Data points represent individual patient samples, with each dot corresponding to a unique analysis region. Analysis performed on singlet T-cells based on hashtag-antibody expression. Statistical comparisons were performed using the Wilcoxon test (paired); p-values are indicated. **(E)** Bar plot showing the distribution of singleton (detected in one cell) and expanded (detected in $\geq 2$ cells) clonotypes across T-cell clusters within different tumor regions per patient. Each bar represents the number of cells from a specific clonotype group in a tumor region in each patient, with colors indicating the relative abundance of distinct T-cell clusters. Analysis performed on singlet T-cells based on hashtag-antibody expression. **(F)** Relative proportions of bystander (light blue), tumor-reactive (light red) and other (gray) T cells across tumor regions. Numbers inside bars represent the absolute number of cells per category. **(G)** Number of detected viral-specific TCRs (against human influenza epitope, HLA-A*02:01-M158–66 (A2+M158)), in each CD8 T cell cluster per tumor-tissue location.

#### Supplemental Tables

**Supplemental Table 1.** Clinical data for all patients included in the study.

**Supplemental Table 2.** List of Differentially Expressed Genes (DEGs) per cluster, related to Figure 1.

**Supplemental Table 3.** Gene signatures from published studies used for the analysis.

**Supplemental Table 4.** List of Differentially Expressed Genes (DEGs) per myeloid and B cell clusters, related to Figure 3.

**Supplemental Table 5.** List of Differentially Expressed Genes (DEGs) per T-cell and NK clusters, related to Figure 4.
